# Identification of TM4SF1 as a novel tumor-associated antigen in biliary tract cancers targetable by CAR-T cell therapy

**DOI:** 10.64898/2026.10.28.762675

**Authors:** Lorraine Nuniz, Julia H. Pham, Juliette Jacques, Corynn Kasap, Jocelin Chen, Jon Akutagawa, Josephine A. Hinneh, Chih-Hao Chang, Cindy Ament, Andrew Lewis, Arun Wiita, Jonathan Chou, R. Katie Kelley, Kwun Wah Wen, Vipul Kumar, Franklin W. Huang

**Author notes:** These authors contributed to the manuscript equally (Lorraine Nuniz, Vipul Kumar). Corresponding Author: Franklin W. Huang, MD PhD, Professor, Division of Hematology/Oncology Department of Medicine, University of California San Francisco, San Francisco Veterans Affairs Medical Center, Address: Medical Sciences Building S1480, 513 Parnassus Ave., San Francisco, CA 94143. **Conflict of interest**. J.C. **and** F.H. are inventors on a patent application related to the CAR T cell technology described in this manuscript (“TM4SF1 CAR CELLS AND METHODS OF USE THEREOF”, U.S. Patent Application No. 63/649,157, U.S. Patent Application No. 63/649,821). The remaining authors declare no competing interests. **Financial Support**. This **study** was funded by the Biohub San Francisco. VK was supported by the ASCO Conquer Cancer Foundation Young Investigator Award. **Author contributions**. Lorraine Nuniz (investigation, validation, formal analysis, methodology supervision, visualization, writing – original draft), Julia H. Pham (investigation, validation, formal analysis), Juliette Jacques (investigation, validation), Corynn Kasap (methodology, resources), Jocelin Chen (resources), Jon Akutagawa (formal analysis, data curation), Josephine Hinneh (methodology, supervision), Chih-Hao Chang (resources), Cindy Ament (resources), Andrew Lewis (Methodology), R. Katie Kelley (resources), Arun Wiita (methodology, resources), Jonathan Chou (methodology, resources), Kwun Wah Wen (methodology, formal analysis), Vipul Kumar (conceptualization, funding acquisition, investigation, validation, formal analysis, project administration, supervision, visualization, writing - original draft, writing - review & editing), Franklin W. Huang (funding acquisition, project administration, supervision, writing - review & editing).

## Abstract

**Background & Aims:** Biliary tract cancers (BTCs) are a rare set of genetically heterogenous and aggressive malignancies associated with delayed presentation and poor prognosis. Thus, there is an unmet clinical need for the development of novel therapeutic strategies. Identification of uniquely expressed cell-surface, tumor-associated antigens (TAAs) holds promise in epithelial tumors more generally, as they can serve as ligands for a variety of therapeutics, including CAR-T cell therapy.

However, TAA identification in BTCs has been limited by expression in normal liver tissue. Here, we aim to characterize the cell-surface protein transmembrane 4 L six family member 1 (TM4SF1) as a potential BTC TAA targetable by CAR-T therapy.

**Methods:** TM4SF1 expression in BTC relative to normal hepatic/biliary tissue was characterized at the mRNA level by analyzing TCGA datasets and at the protein level by immunohistochemistry of archival primary BTC tissue. Anti-TM4SF1 CAR-T cells were evaluated through *in vitro* coculture assays against human-derived BTC cell lines and in a cell line-derived xenograft (CDX) model.

**Results:** We find TM4SF1 is significantly upregulated in BTC relative to normal hepatic/biliary tissue and across BTC anatomical subtypes at the mRNA and protein level. To validate TM4SF1 as a targetable TAA in BTC, we show anti-TM4SF1 CAR-T cells demonstrate robust growth inhibition of BTC cell lines *in vitro* and significant anti-tumor activity against a CDX *in vivo*.

**Conclusions:** Together, our data provides support for TM4SF1 as a promising TAA in BTCs that can be used for the rational development of targeted therapeutic modalities, including CAR-T cells, in a patient population with significant unmet need.

**Impact & Implications:** TM4SF1 has been linked across various solid tumors as a driver of disease aggressiveness and poor outcomes. Our work seeks to address the need for more effective BTC therapies by investigating TM4SF1 as a potential target. Our results establish TM4SF1 as a potential TAA in BTC targetable by CAR-T therapy and support further preclinical investigation into anti-TM4SF1 targeted therapies in BTC.

**Highlights:**

- TM4SF1 is upregulated in BTC relative to normal tissue at the mRNA and protein level.
- Anti-TM4SF1 CAR-T cells demonstrate strong, dose-dependent growth inhibition *in vitro*
- Anti-TM4SF1 CAR-T cells demonstrate robust tumor control *in vivo*

## Introduction

Biliary tract cancers (BTCs) comprise a rare but aggressive, heterogeneous group of invasive neoplasms stemming from the biliary system. They are grouped by anatomic site of origin and include intrahepatic cholangiocarcinoma (IHCC), extrahepatic cholangiocarcinoma (EHCC), and gallbladder carcinoma (GBC)^1^. The current standard of care treatment for patients with advanced BTCs is cisplatin and gemcitabine combined with immune checkpoint inhibition with either durvalumab or pembrolizumab^2,3^, but curative treatment remains elusive. One of the many challenges in obtaining durable responses is the genetic heterogeneity between IHCC, EHCC, and GBC^1^, despite a number of actionable targets being identified. For example, IDH1, BRAF, and FGFR2 alterations are found in a subset of IHCC, while KRAS mutations and HER2 amplification are associated with a subset of EHCC^1,4–6^. Moreover, despite the availability of IDH, BRAF, and FGFR-targeted therapies in biomarker-selected populations after frontline therapy, durable responses are limited, and most patients succumb to their illness with a median overall survival of approximately one year^2,3^. Thus, there is an urgent, unmet clinical need to identify novel targets and therapies for patients with advanced BTCs.

Targeting tumor-specific cell-surface proteins, known as tumor-associated antigens (TAAs), holds great promise for the treatment of epithelial and other tumors. Identification of TAAs is translationally impactful as they can serve as ligands for a variety of clinical therapeutics, ranging from radioligand to CAR-T cell therapy. However, identification of such targets has been met with limited success, owing to promiscuous expression of potential TAAs in normal tissues. TAA development in BTCs has been further hampered by expression in normal liver tissue, leading to dose-limiting off-tumor, on-target hepatotoxicity^7^. TAAs such as Claudin 18.2 and HER2 are currently in development; however, limited expression across BTC subtypes, amongst other factors, limits their widespread use, thus leaving an unmet need to identify novel BTC TAAs^7,8^.

Transmembrane-4 L-six family member-1 (TM4SF1) is a small plasma membrane glycoprotein and regulator of cell adhesion, motility, proliferation, and angiogenesis. In a disease context, TM4SF1 has been implicated in increased tumor growth, invasion, and metastasis^9^, and pre-clinical studies have demonstrated its upregulation in both tumor cells and endothelial cells lining tumor vasculature^10,11^. Clinically, high TM4SF1 expression is associated with more aggressive disease and poor prognosis across a variety of carcinomas, including bladder, pancreatic, hepatocellular, colorectal, and ovarian cancers ^9,12^, leading to interest in TM4SF1 as a potential therapeutic target. Correspondingly, preclinical studies have shown feasibility in targeting TM4SF1 by several modalities, including monoclonal antibodies (mAb) in multiple solid tumors^13–16^ and CAR-T cells in ovarian cancer and urothelial carcinoma cell lines^17,18^. Additionally, a clinical stage TM4SF1-targeting antibody-drug conjugate is in development evaluating activity in multiple solid tumors (NCT06440005). Collectively, the current body of data provides a strong rationale that TM4SF1 could serve as an actionable TAA across a diversity of epithelial tumors. However, whether TM4SF1 might serve as a TAA and therapeutic target in BTCs remains unknown.

Here, we investigated TM4SF1 as a potential TAA in BTCs through comprehensive expression analysis of primary patient tissue and human-derived BTC cell lines at the mRNA and protein levels as well as through pre-clinical experiments testing the *in vitro* and *in vivo* efficiency of anti-TM4SF1 CAR-T cells against human-derived BTC cell lines and xenograft models.

## Methods

### TCGA analysis of TM4SF1 expression

RSEM normalized counts of TM4SF1 from TCGA were downloaded from the <u>UCSC</u> <u>Xena browser</u>^19^. These datasets were reanalyzed and harmonized to eliminate computational batch effects. 9,184 TCGA samples highly expressed TM4SF1, with log2(RSEM normalized count +1) > 5. Of these, 36 were cholangiocarcinoma tumor samples and 9 were matched normal samples. Box and violin plots were generated with Python (v.3.11.13) packages, seaborn (v0.11.2), and matplotlib (3.8.1). Mann-Whitney U tests were performed to determine significant differences between median expected counts.

### Immunohistochemistry

Immunohistochemistry (IHC) protein expression analysis was performed for all samples using a previously validated anti-human TM4SF1 antibody (Abcam, ab113504) as previously described^18^. De-identified patient samples were obtained internally at UCSF as part of an IRB-approved tissue banking protocol (IRB #12-09576) from archival excision specimens of patients with IHCC (n=17 samples from x=9 patients), EHCC (n=6 samples from x=3 patients), and GBC (n=6 samples from x=5 patients) with matched normal samples, where available. A cholangiocarcinoma tissue microarray (TMA) was obtained commercially (TissueArray.com, GA802b). Slides were reviewed by a board-certified GI pathologist and imaged on the Olympus BX 53 microscope with the Olympus DP73 camera and Olympus cellSens Entry 4.3 (Build 31056) acquisition software. For UCSF samples, the percent of tumor or normal epithelial cells with significant staining for TM4SF1 was scored (“percent positivity”); for the TMA, cores were similarly scored for percent positivity and then placed into categories defined as diffuse (>90%), patchy (10-90%), focal (<10%), or negative (0%).

### Flow cytometry

Flow cytometry was performed to evaluate cell-surface protein expression of TM4SF1 across a panel of human-derived BTC cell lines including NOZ, RBE, HuCCT-1, and SNU-308 as previously described for urothelial carcinoma cell lines^18^. Briefly, cells were stained with an anti-TM4SF1-APC antibody (Miltenyi, clone REA851, 1:100) alongside a mouse IgG1 κ isotype control (BD Biosciences, clone MOPC-21, 1:100), and median fluorescent intensity (MFI) shift relative to the isotype control was calculated as a measure of cell-surface expression (FlowJo version 10.10.0)

### TM4SF1 CAR-T cell generation

Anti-TM4SF1 CAR-T cells were generated as previously described^18^, with the scFv in the VH-VL configuration, as were anti-BCMA CAR-T cells as a specificity control^20^. Briefly, primary human T-cells from two separate, healthy donors were isolated from peripheral blood leukopaks. Concentrated lentivirus and ImmunoCult™ Human CD3/CD28 T Cell Activator (Stemcell Technologies, 10971) were added, and following 72hr incubation, virus and activator were removed. Cells were subsequently expanded and used between day 12-16 for downstream assays, during which flow cytometry was performed to evaluate transduction efficiency via GFP expression. Cells were grown in TexMACS™ Medium (Miltenyi Biotech, 130-097-196) supplemented with human recombinant cytokines IL-7 and IL-15 added at 10ng/mL (Stemcell Technologies, 78053.1; 78031.1).

### Cell culture

The established human-derived BTC cell lines HuCCT-1 (56M, IHCC), SNU-308 (unknown, GBC), NOZ (48F, GBC), and RBE (64F, IHCC) were obtained as kind gifts from Dr. John Gordan (UCSF). Cells were grown in standard RPMI media (Gibco) supplemented with 10% FBS (Cell Culture Collective) and 1% penicillin/streptomycin. All experiments were conducted within 20 passages from the parental stock. Cell lines were routinely tested for mycoplasma (Lonza).

### IncuCyte co-culture assays

Co-culture assays were performed as previously described for urothelial carcinoma cell lines^18^ with BTC cell lines transduced with NucLight Red virus (NLR; Sartorius). Briefly, cell lines were plated at 5k cells/well in a 96-well plate and allowed to adhere overnight; T-cells were either thawed and allowed to recover overnight or used fresh. The following day, anti-TM4SF1 CAR-T cells or control untransduced (UTD), empty vector, or anti-BCMA CAR-T cells were added to cancer cells over a range of effector-to-target (E:T) ratios. Target cell killing was monitored through IncuCyte S3 (Sartorius) live-cell imaging, with killing quantified through red object count per well normalized to d0h0m0 values. All data was plotted on Prism (GraphPad, v10).

### Animal studies

All animal studies were performed under an approved UCSF Institutional Animal and Use Committee (IACUC) protocol (AN202104-00E) as previously done for urothelial xenografts18 with the following modifications. HuCCT-1 cells were subcutaneously implanted at 3×10^6^ cells per tumor in 8-10wk old NSG male mice (d0). When tumor volumes reached between 50-100 mm^3^ (d19), mice were randomized into two treatment groups and treated with 3-5×10^6^ UTD T-cells or anti-TM4SF1 CAR-T cells. Body weight (BW) and tumor volume (TV) measurements were performed by personnel from the UCSF Preclinical Therapeutics Core in a blinded fashion twice weekly, then once weekly after d74. Two UTD-treated mice were taken down early (d84) due to tumor ulceration; all remaining mice were euthanized on d105 for tumor-unrelated reasons due to spread of *Corynebacterium bovis* infection. For tissue analysis, primary tumors were collected, fixed in paraffin, and stained for IHC as described above; comparisons in tumor cell density and IHC staining between control-treated and CAR-T-treated mice were made by a pathologist to confirm whether tissue was genuine tumor or residual mass and to establish potential changes in TM4SF1 protein expression.

### Statistics and reproducibility

Sample sizes for CAR-T experimental cohorts were determined based on expected effect size and threshold for achieving biological replicants. Data from IHC staining and *in vitro* co-culture assays are shown as mean□±□standard error of the mean (SEM). Statistical significance for all tests was set at p□<□0.05.

## Results

### TM4SF1 is upregulated at the mRNA and protein level in BTCs relative to normal tissue

We first evaluated TM4SF1 mRNA expression across all tumors available in the TCGA database and found BTCs had the second-highest expression (Figure 1A), which raised the initial possibility of TM4SF1 serving as a TAA in BTCs. We then performed a focused analysis comparing TM4SF1 mRNA expression in matched BTC-normal tissue from the TCGA-CHOL dataset and found significantly higher expression in BTC tissue (median expression 14.35 vs 11.53, p=1.353e-4, Mann-Whitney U) (Figure 1B).

**Fig. 1.**
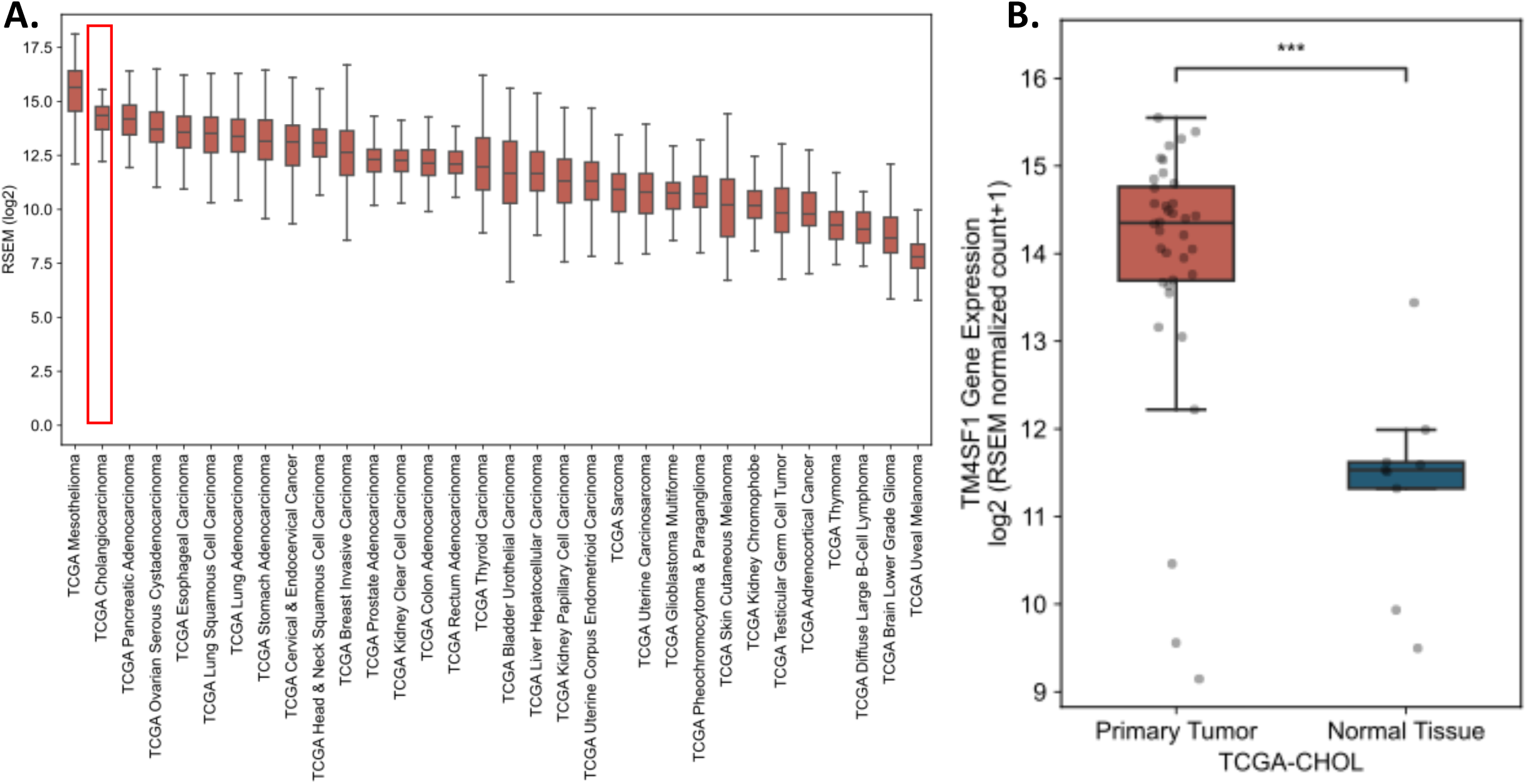
TM4SF1 mRNA upregulation in biliary tract cancers. **(A)** TM4SF1 mRNA expression across all tumor types in the TCGA database, with cholangiocarcinoma (red box) showing the second highest TM4SF1 expression . **(B)** TM4SF1 mRNA expression in cholangiocarcinoma (TCGA-CHOL n=36) compared to normal liver tissue (TCGA-CHOL n=8) (median expression 14.35 vs 11.53; ***, p=1.353e-4, Mann-Whitney U).

To further assess TM4SF1 upregulation at the protein level, we performed immunohistochemistry (IHC) protein expression analysis of TM4SF1 across BTC anatomical subtypes in a cohort of primary patient samples at our institution (n=29 unique samples across x=17 unique patients) including IHCC, EHCC, and GBC samples and in a commercial cholangiocarcinoma tissue microarray (TMA) including IHCC and EHCC cores, with IHC analysis performed using a previously validated anti-TM4SF1 antibody^18,21–22^. Across our cohort of patient samples, we found TM4SF1 protein expression across the three BTC subtypes was significantly increased ∼15.8-fold in BTC tissue relative to normal hepatobiliary parenchymal tissue (73.33% vs 4.64% percent positivity, P<0.0001) (Figure 2A-C). In our cholangiocarcinoma TMA, we similarly confirmed strong TM4SF1 protein expression across the majority of cores, with 83.33% of cores (39/45 IHCC and 16/21 EHCC cores) showing diffuse staining (>90% positivity), 12.12% (5/45 IHCC and 3/21 EHCC cores) showing patchy staining (10-90% positivity), 3.03% (2/21 EHCC cores) showing weak staining (<10% positivity) and only 1.52% (1/45 IHCC cores) showing negative staining (0% positivity) (Figure 2D-E, Figure S1).

**Fig. 2.**
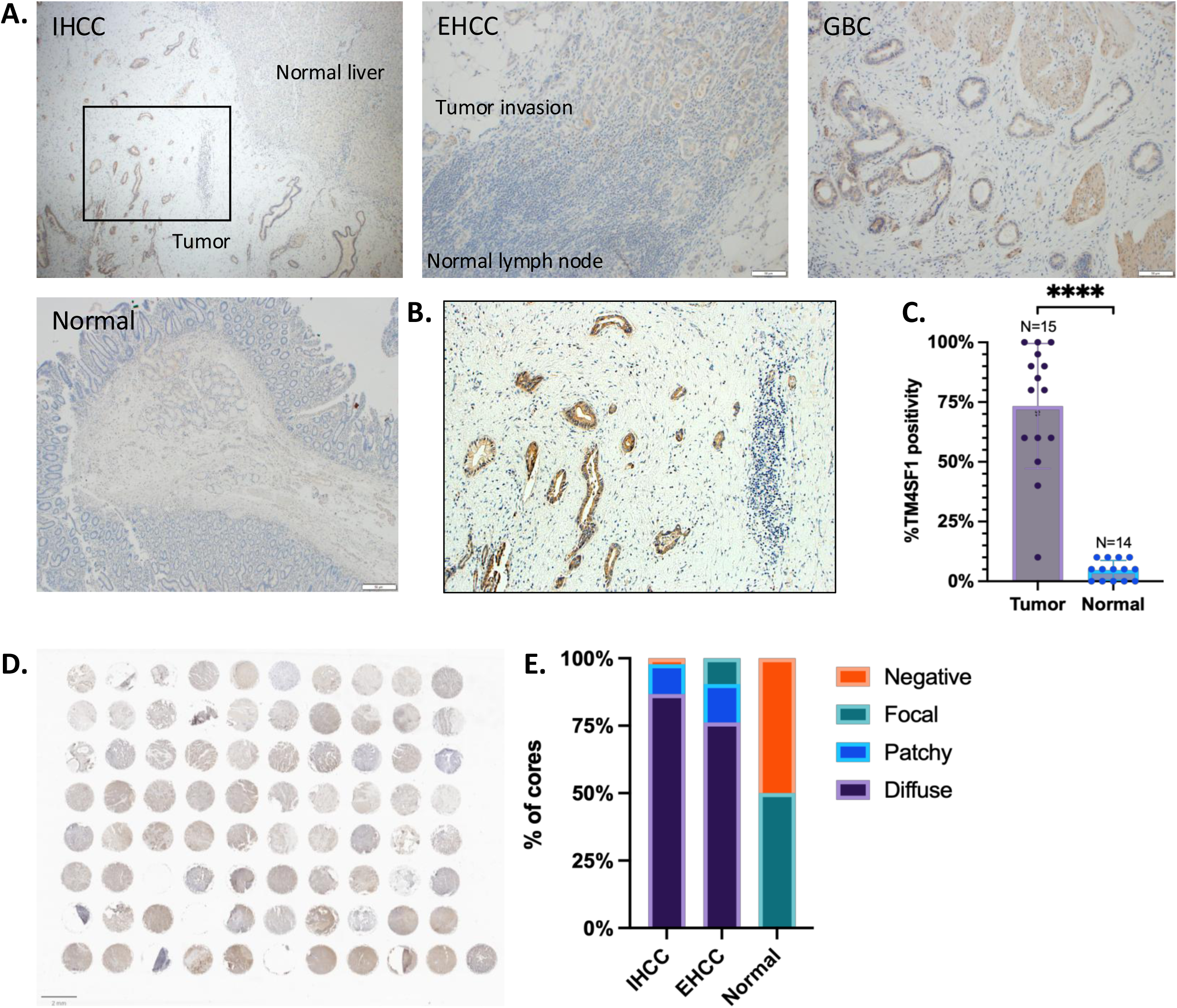
TM4SF1 protein upregulation across subtypes of biliary tract cancer (BTC). **(A)** Representative images of TM4SF1 IHC staining across BTC anatomical subtypes, showing TM4SF1 expression in primary intrahepatic cholangiocarcinoma with adjacent normal liver tissue ("IHCC", 40X), extrahepatic (distal) cholangiocarcinoma lymph node invasion ("EHCC", 100X), primary gallbladder carcinoma tissue ("GBC", 100X), and adjacent normal duodenum from an EHCC patient ("Normal", 40X). **(B)** Blow-up of TM4SF1 expression in IHCC tissue from (A) (100X). **(C)** Summary of TM4SF1 protein expression across BTC anatomical subtypes from matched tumor-normal samples (n=12), unmatched tumor samples (n=3), unmatched normal liver/bile duct samples (n=2) by IHC analysis, with TM4SF1% positivity (percent of epithelial cells with real TM4SF1 staining) scored by GI pathologist (P<0.0001). **(D)** TM4SF1 protein expression across cholangiocarcinoma tissue microarray (US BiomaxGA802b), with **(E)** expression in IHCC cores (n=45), EHCC cores (n=21), and normal liver cores (n=4) measured by IHC; sample cores scored by GI pathologist for percent of tumor cells with real TM4SF1 staining as diffuse (>90%), patchy (10-90%), focal (<10%), or negative (0%).

To validate TM4SF cell-surface protein expression, we performed flow cytometry analysis of a panel of four human-derived BTC cell lines, including two IHCC cell lines (RBE, HuCCT-1) and two GBC cell lines (NOZ, SNU-308). Surface expression was measured through median fluorescence intensity (MFI) shift of TM4SF1-stained cells relative to a mouse IgG1 κ isotype control, with all four cell lines showing medium to high TM4SF1 cell-surface expression (Figure 3A).

**Fig. 3.**
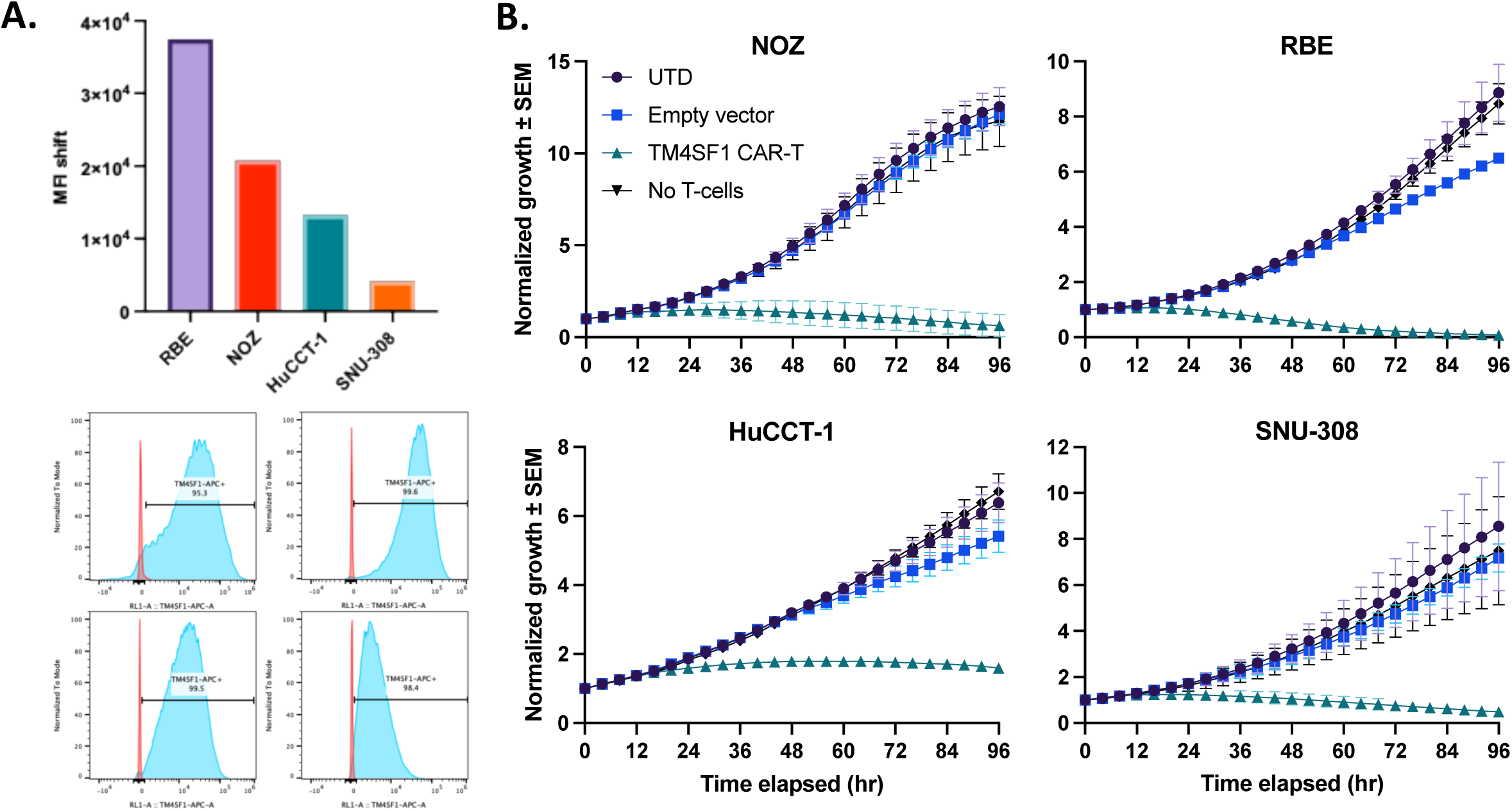
Anti-TM4SF1 CAR-T cells induce growth inhibition of BTC cell lines *in vitro*. **(A)** Flow cytometry analysis of TM4SF1 cell-surface expression across a panel of human-derived BTC cell lines NOZ, RBE, HuCCT-1, and SNU-308, with TM4SF1-APC shift (cyan) shown relative to Mouse IgG1,κ APC isotype control (red); TM4SF1 surface expression quantified through median fluorescence intensity (MFI) shift of TM4SF1 relative to isotype control . **(B)** *In vitro* growth inhibition of BTC cell lines (Nuclight Red) by anti-TM4SF1 CAR-T cells, with cell growth over 96hr duration quantified through red object count per well normalized to d0h0m0; mean ± SEM shown (n=3 biological replicates, each with n=3 technical replicates). Activity of anti-TM4SF1 CAR-T cells compared to control untransduced (UTD) T-cells, empty vector T-cells, and baseline growth without added T-cells, with cocultures performed at effector-to-target (E:T) ratio of 1:1.

### Anti-TM4SF1 CAR-T cells demonstrate robust killing of BTC cell lines *in vitro* and ***in vivo***

Having confirmed TM4SF1 surface expression in BTC cell lines, we tested the pre-clinical efficacy of our previously developed, second-generation anti-TM4SF1 CAR-T cells^18^ for their ability to inhibit BTC cell line growth. In *in vitro* co-culture assays, we found anti-TM4SF1 CAR T-cells derived from two independent donors demonstrated potent, dose-dependent growth inhibition of all four cell lines compared to both control untransduced (UTD) T-cells and empty vector T-cells transduced with an scFv-less CAR construct (E:T 1:1, Donor 1 shown in Figure 3B; full dose range from both donors shown in Figure S2). We additionally generated and tested anti-BCMA CAR-T cells, which target a non-BTC, plasma-cell lineage marker^23^, as a specificity control and found no significant off-target growth inhibition when compared to treatment with anti-TM4SF1 CAR-T cells (Figure S3).

Next, we evaluated the efficacy of anti-TM4SF1 CAR-T cells *in vivo* using a heterotopic cell-line derived xenograft (CDX) model with the cell line HuCCT-1, which has previously been utilized in pre-clinical xenograft experiments^24^. Treatment of mice with anti-TM4SF1 CAR-T cells resulted in a 100% response rate, with 5/8 mice experiencing durable, complete tumor regression without palpable tumors over 100 days (“complete response”) and 3/8 mice experiencing a significant partial response, with an initial 63.1% decrease in tumor volume followed by eventual tumor progression (“partial response”) (Figure 4A). Despite progression, the overall mean tumor volumes during the follow-up period of the partial-response group remained significantly reduced in comparison to the control UTD-treated group (269.95mm^3^ vs 859.67mm^3^ on d105, p=0.0033, one-way ANOVA), the latter in which all 8/8 mice experienced continuous progressive tumor growth, with 2/8 mice reaching humane endpoints by day 84. To investigate antigen downregulation as a potential mechanism of tumor recurrence in the partial-response mice, we conducted IHC analysis of primary tumors from CAR-T-treated partial responders compared to control-treated mice but observed no difference in TM4SF1 protein expression nor tumor cell density (Figure 4B).

**Fig. 4.**
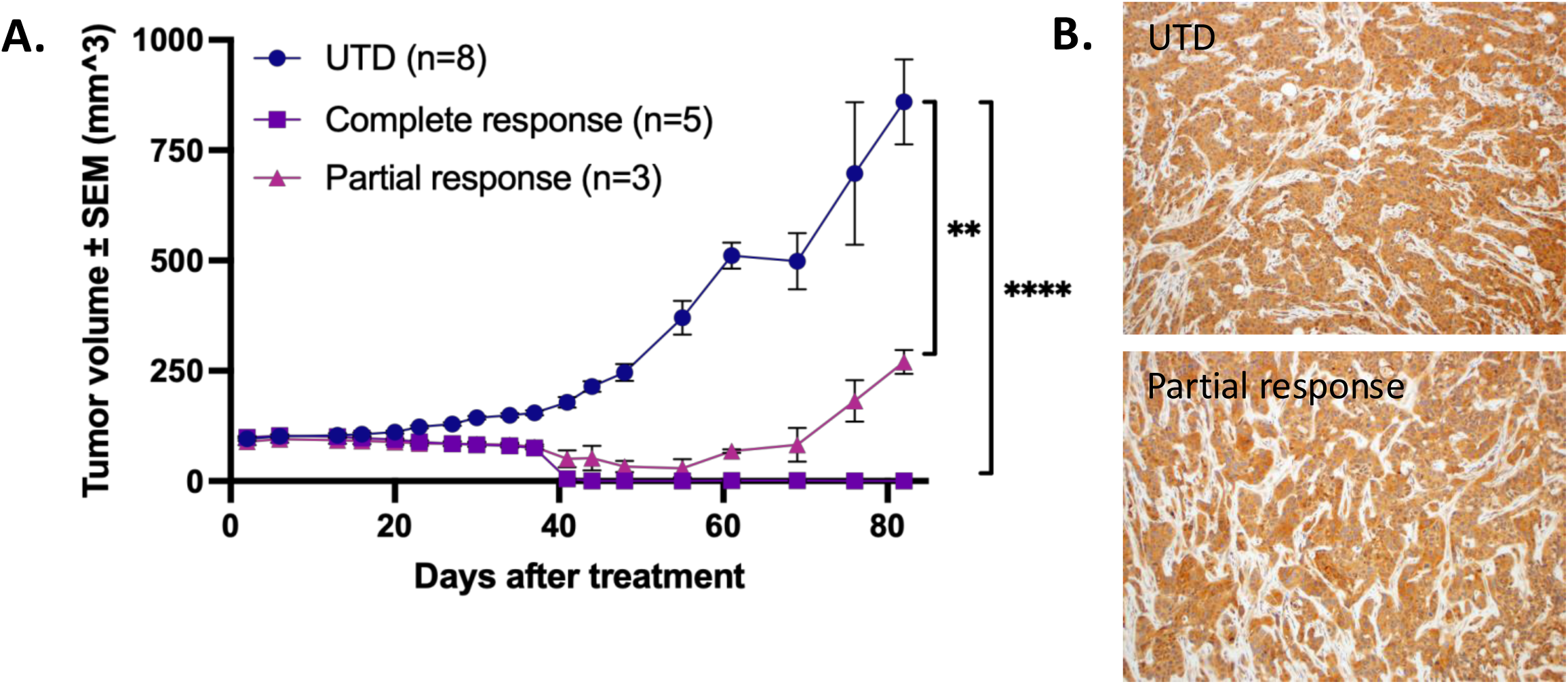
Anti-TM4SF1 CAR-T cells demonstrate robust tumor control *in vivo*. **(A)** Tumor growth ± SEM following treatment of mice with control untransduced T-cells (“UTD”) or anti-TM4SF1 CAR-T cells (n=8 per group). All CAR-T-treated mice showed significant reductions in tumor volumes relative to control-treated mice, with 5/8 treated mice showing complete tumor regression (“complete response”; **** P<0.0001) and 3/8 treated mice showing partial tumor regression followed by recurrence (“partial response”;** P=0.0011). P values were determined using one-way ANOVA followed by Tukey multiple comparison test. **(B)** Absence of TM4SF1 downregulation in recurrent tumors of "partial response" mice relative to UTD-treated mice via TM4SF1 immunohistochemistry (100X).

Collectively, our results show that anti-TM4SF1 CAR-T cells provide robust, dose-dependent growth inhibition of BTC cell lines *in vitro* and significant tumor control of HuCCT-1-derived xenografts *in vivo*.

## Discussion and Conclusions

Biliary tract cancers are a family of disease marked by aggressive, heterogenous tumors and an absence of effective therapeutic interventions, resulting in devastating patient outcomes. In this study, we sought to identify additional TAAs capable of serving as novel therapeutic targets in BTCs to address this unmet clinical need.

In our expression analysis, we found marked TM4SF1 upregulation in BTCs relative to normal hepatobiliary tissue at the mRNA and protein level, the latter revealing even more profound upregulation based on our cohort; while limited in sample number and hence our ability to make quantitative cross-subtype comparisons, the majority of tumor samples for a majority of patients showed strong TM4SF1 protein expression. To further validate TM4SF1 as a novel TAA for BTCs, we subsequently evaluated our previously designed anti-TM4SF1 CAR-T cells against a panel of human-derived BTC cell lines in both *in vitro* and *in vivo* models. Having confirmed TM4SF1 surface expression via flow cytometry in our four cell lines, we demonstrated potent, dose-dependent growth inhibition of all cell lines in *in vitro* coculture assays as well as robust tumor control in a heterotopic xenograft model, with all CAR-T-treated mice showing a significant response compared to control-treated mice. In this context, while a minority of mice did show partial response characterized by initial tumor regression followed by recurrence, tumor size remained significantly smaller than control-treated mice and differences in TM4SF1 expression were not observed, suggesting therapeutic escape mechanisms other than antigen downregulation, as consistent with observations in a separate ovarian cancer CDX model of anti-TM4SF1 CAR-T cells^17^.

Prior efforts to identify TAAs in BTCs and other epithelial tumors have been limited by promiscuous expression in normal tissue, both within the parental tissue of cancer origin as well as across other tissues. While we acknowledge that TM4SF1 has some degree of expression in several other normal tissues, including lung^18^, this is also the case with other clinical stage TAAs in other epithelial tumors, such as STEAP1 and PSMA in prostate cancer^25^ and EGFR in lung and colorectal cancers^26,27^. Ultimately, definite toxicity testing will have to advance to assess potential clinical impacts; in this context, ongoing Phase 1 clinical trials of the anti-TM4SF1 ADC AGX101^28^ may be informative in this regard. However, it is reassuring that we find minimal, if any, expression in normal hepatobiliary tissue in our cohort.

Beyond promiscuous expression, the targeting of TAAs in BTCs is frequently hindered by antigenic heterogeneity as well as BTCs’ characteristically immunosuppressive microenvironment^29^. While our utilization of an immunodeficient, cell line-derived model limits our ability to directly interrogate the efficacy of our anti-TM4SF1 CAR-T cells in face of these challenges, the observed TM4SF1 protein expression patterns and robust tumor control *in vivo* are nonetheless promising initial results which add to a growing body of evidence supporting TM4SF1 as a potential therapeutic target in disease targetable by multiple modalities which merits further investigation. We also note that beyond CAR-T cell monotherapy, therapeutic strategies incorporating TM4SF1, including logic-gated approaches with TM4SF1 serving as one of several specificity ligands^30^ and combination (immuno)therapies, may be considered to address the host of challenges faced by BTC TAAs. In summary, our study represents a preclinical advance in the field of BTC therapeutic development by nominating TM4SF1 as a potentially relevant TAA targetable by CAR-T cell therapy.

## Abbreviations

ADC: antibody-drug conjugate
APC: allophycocyanin
BCMA: B-cell maturation antigen
BRAF: B-Raf proto-oncogene, serine/threonine kinase
BTC: biliary tract cancer
CAR-T: chimeric antigen receptor T-cell therapy
CDX: cell line-derived xenograft
EHCC: extrahepatic cholangiocarcinoma
FGFR2: fibroblast growth factor receptor 2
GBC: gallbladder carcinoma
GFP: green fluorescent protein
HER2: human epidermal growth factor receptor 2
IACUC: Institutional Animal Care and Use Committee
IDH1: isocitrate dehydrogenase 1
IHC: immunohistochemistry
IHCC: intrahepatic cholangiocarcinoma
KRAS: Kirsten rat sarcoma viral oncogene homolog
mAb: monoclonal antibody
MFI: median fluorescent intensity
NLR: NucLight Red (Sartorious)
NSG: NOD scid gamma
PSMA: prostate-specific membrane antigen
SEM: Standard Error of the Mean
ScFv: single-chain variable fragment
STEAP1: six transmembrane epithelial antigen of the prostate 1
TAA: tumor-associated antigen
TCGA: The Cancer Genome Atlas
TMA: tissue microarray
TM4SF1: transmembrane 4 L six family member 1
UTD: untransduced

## Acknowledgements

This project was supported by the Biohub San Francisco (F.W.H.). V.K was supported by an ASCO Conquer Cancer Foundation Young Investigator Award. We thank the Helen Diller Family Comprehensive Cancer Center’s Preclinical Therapeutics Core at UCSF for experimental design & technical support. We thank Elizabeth Yip and Henry Delavan for xenograft model experimental design & technical assistance. We thank Scarlett Wen for technical assistance with patient samples. We thank all members of the Huang lab for their contributions.

