## Supplemental figures 1-3 for "Identification of TM4SF1 as a novel tumor-associated antigen in biliary tract cancers targetable by CAR-T cell therapy"

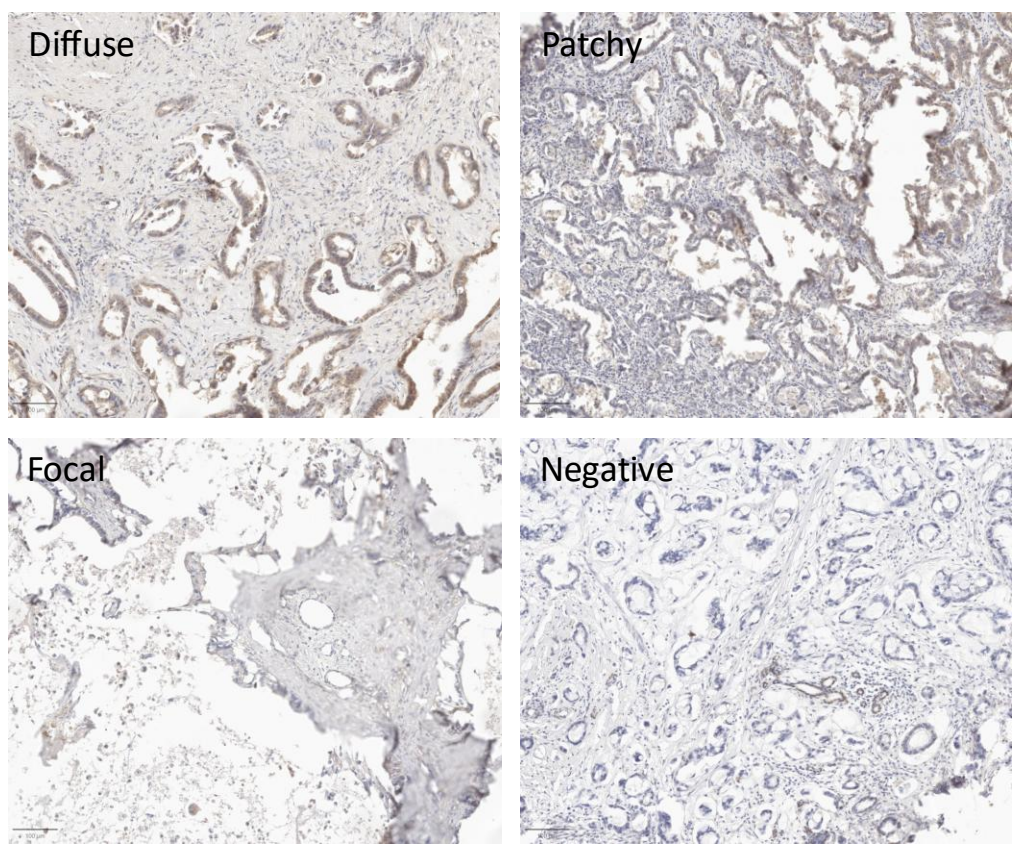

**Supplementary Fig. 1. TM4SF1 protein expression across cholangiocarcinoma TMA.** Representative cores showing diffuse (top left), patchy (top right), focal (bottom left), and negative (bottom right) IHC staining for TM4SF1 at 10X magnification.

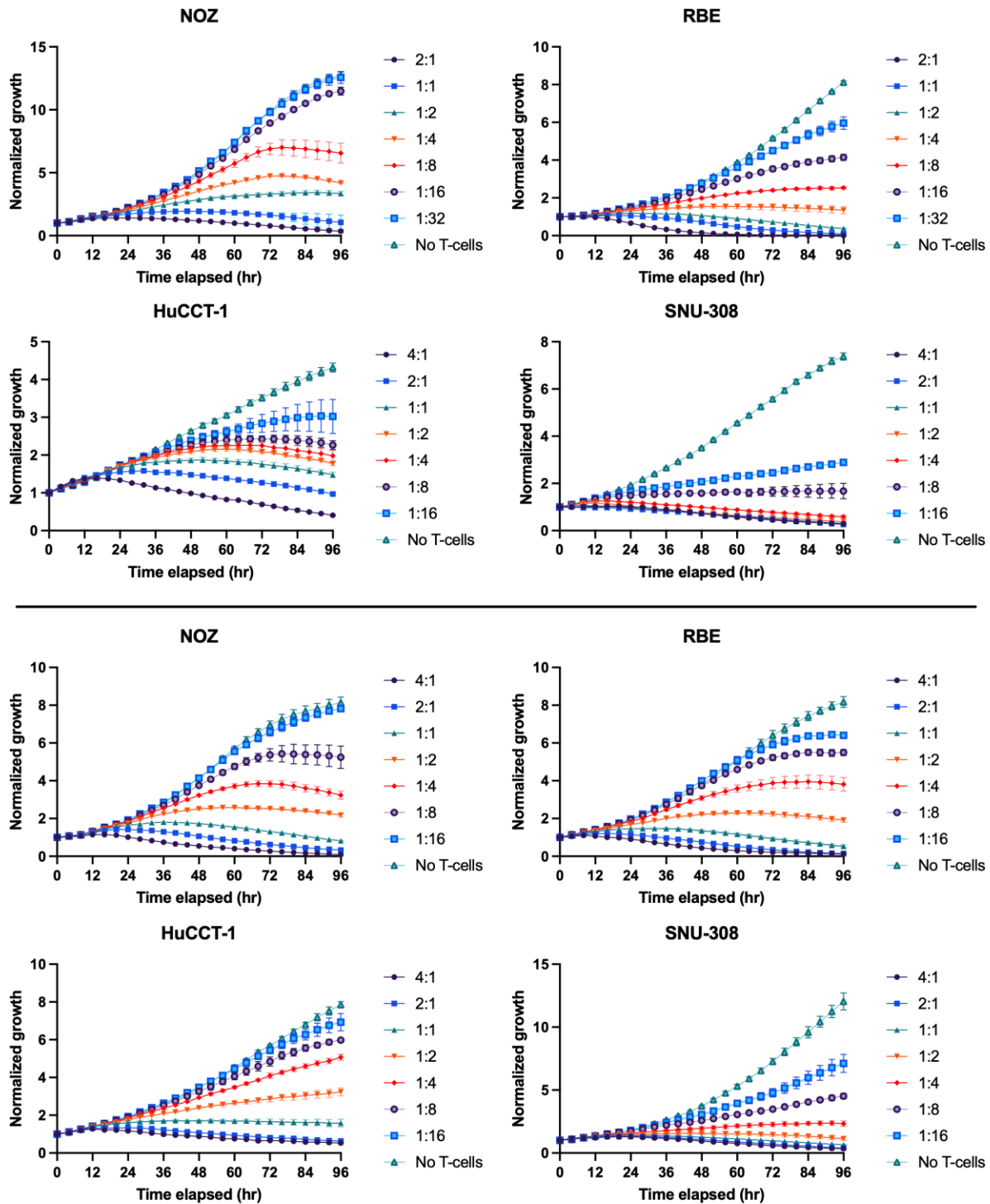

**Supplementary Fig. 2. Dose-dependent responsiveness of anti-TM4SF1 CAR-T cells across donors.** Dose-dependent response of anti-TM4SF1 CAR-T cells across two separate T-cell donors (top: Donor 1, bottom: Donor 2) across effector-to-target (E:T) ratios from 4:1 to 1:16, 1:32 across BTC cell lines NOZ, RBE, HuCCT-1, and SNU-308 compared to baseline growth with no added T cells.

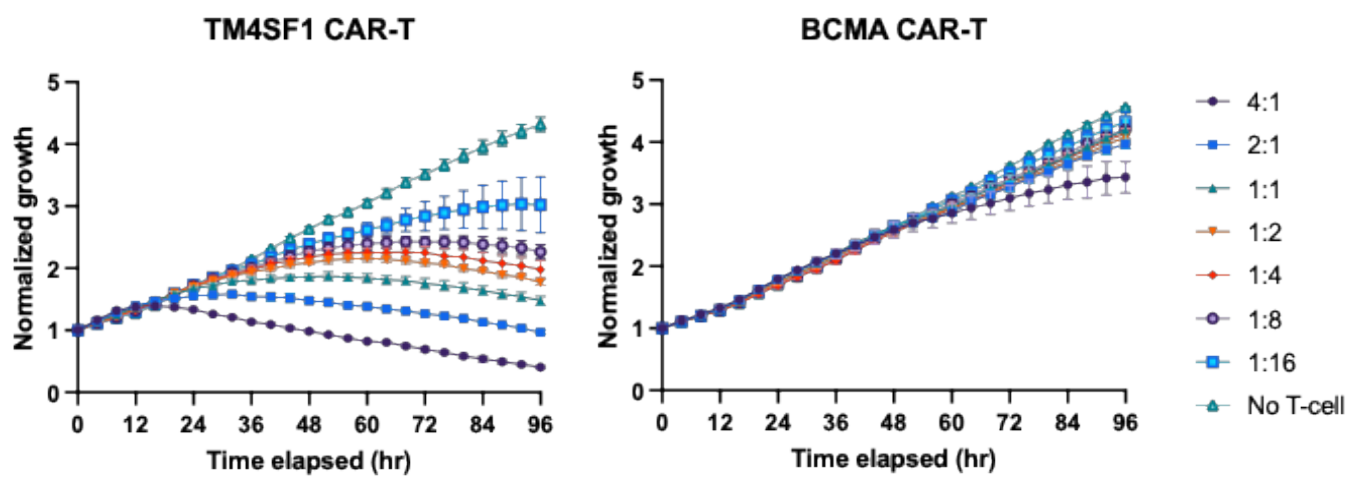

**Supplementary Fig. 3. Response to anti-TM4SF1 versus anti-BCMA CAR-T specificity control.** *In vitro* growth inhibition by anti-TM4SF1 CAR-T cells compared to absence of growth inhibition by anti-BCMA CAR-T cells (targeting non-BTC, plasma-cell lineage marker) demonstrates target-specific cytotoxicity of CAR construct; shown at E:T ratio of 4:1 to 1:16 for representative BTC cell line HuCCT-1 . Similar findings in other BTC cell lines not shown.
